# Determination of hemicellulose, cellulose, holocellulose and lignin content using FTIR in *Calycophyllum spruceanum* (Benth.) K.Schum. and *Guazuma crinita* Lam.

**DOI:** 10.1101/2021.09.01.458618

**Authors:** Rosario Javier-Astete, Jorge Jimenez-Davalos, Gaston Zolla

**Affiliations:** Grupo de Investigacion en Mutaciones y Biotecnologia Vegetal, Facultad de Agronomia, Universidad Nacional Agraria La Molina, Lima, Peru; Laboratorio de Fisiologia vegetal, Facultad de Ciencias, Universidad Nacional Agraria La Molina, Lima, Peru

**Author notes:** Corresponding author: (GZ).

**Keywords:** Capirona, Bolaina, FTIR spectroscopy, PLS, chemometrics

## Abstract

Capirona (*Calycophyllum spruceanum* (Benth.) K. Schum.) and Bolaina (*Guazuma crinita* Lam.) are fast-growing Amazonian trees with increasing demand in timber industry. Therefore, it is necessary to determine the content of cellulose, hemicellulose, holocellulose and lignin in juvenile tress to accelerate forest breeding programs. The aim of this study was to identify chemical differences between apical and basal stem of Capirona and Bolaina to develop models for estimating the chemical composition using Fourier transform infrared (FTIR) spectra. FTIR-ATR spectra were obtained from 150 samples for each specie that were 1.8 year-old. The results showed significant differences between the apical and basal stem for each species in terms of cellulose, hemicellulose, holocellulose and lignin content. This variability was useful to build partial least squares (PLS) models from the FTIR spectra and they were evaluated by root mean squared error of predictions (RMSEP) and ratio of performance to deviation (RPD). Lignin content was efficiently predicted in Capirona (RMSEP = 0.48, RPD > 2) and Bolaina (RMSEP = 0.81, RPD > 2). In Capirona, the predictive power of cellulose, hemicellulose and holocellulose models (0.68 < RMSEP < 2.06, 1.60 < RPD < 1.96) were high enough to predict wood chemical composition. In Bolaina, model for cellulose attained an excellent predictive power (RMSEP = 1.82, RPD = 6.14) while models for hemicellulose and holocellulose attained a good predictive power (RPD > 2.0). This study showed that FTIR-ATR together with PLS is a reliable method to determine the wood chemical composition in juvenile trees of Capirona and Bolaina.

## Introduction

The Amazonian Forest plays an important role in human being and environmental welfare; however, the growing deforestation could reach a summit point thus this forest will turn to non-forest ecosystem [1,2]. Just in the last 10 years, agriculture, mining, narcotraffic and growing population [2] have contributed to the disappearance of 26 million Ha of Amazonian Forest [3], an area close to New Zealand surface. Thus, Amazonian deforestation has increased the intensity of global warming, loss of biodiversity and affects indigenous communities adversely (migration, prostitution, etc) [2]. Deforestation also increases risk of human contact with zoonotic pathogens [4].

In this context, tree breeding is a competitive and environmentally friendly opportunity to recover and increase woody biomass [5]. Tree breading programs increase wood quality and forestry production; such is the case of *Scots pine*, *Norway spruce*, *Stika spruce*, *Eucalyptus pellita* and *Acacia mangium* [5,6]. In the amazon forest there is interest in tree breading programs of Capirona and Bolaina [7]. These fast-growing trees deliver wood products at an early stage [8] and can be coppiced for successive harvests [9]. Both species have a high wood market demand and are widely used for reforestation [10–12]. Furthermore, the production of both species is a sustainable activity for small farmers across Amazon and provides environmental services like carbon capture, conservation of biodiversity and soil fertility [8].

Selection of plus-tree is normally based on visual properties rather than quality parameters such as cellulose, hemicellulose and lignin content [5,6]. The chemical composition is a key feature that determines wood quality and the end-use of wood in early stage [13,14]. For instance, lignin is particularly appreciated in timber industry but not in pulp production, while cellulose is quite valued in pulp industry [13]. The information of chemical composition can help breeders in the selection process of plus-trees [6]. However, determination of chemical composition using conventional methods is expensive (USD 1000 per tree), time consuming, destructive and requires chemical reagents [6,15].

On the contrary to conventional techniques, FTIR-ATR spectroscopy is a practical and economical analytic method to determine wood chemical composition [15]. This method has been successfully used to identify cellulose, hemicellulose, lignin, monosaccharide, extractive compounds and proteins in forest species [15–18]. The complex data generated by FTIR-ATR (vibration of chemical bind and functional groups) is processed by multivariate analysis [19] such as partial least squared (PLS). This multivariate method is one of the most used ones and can predict sample properties by a number of latent variables which compresses spectra information [20]. Despite the successful application and numerous advantages of these methods, they are barely applied in native Amazonian trees.

To the best of our knowledge, there is no FTIR-ATR spectroscopy coupled to PLS in fast growing trees in juvenile stages. Therefore, the aim of this work was to develop efficient models to predict cellulose, hecmicellulose, holocellulose and lignin content in juvenil trees of Capirona and Bolaina.

## Material and Methods

### Plant Material

100 Capirona trees and 150 Bolaina trees acquired from Ayahuasca Ecolodge E.I.R.L. and Est. Exp. Agraria Pucallpa – Ucayali – INIA, were kept in a net house in La Molina (12°05’ S, 76°57’ W and 243.7 masl). When trees reached 1.8 year-old, the samples were collected from apical and basal stem to expand range of wood chemical variability. We grouped 4 and 6 plants as a sample for Capirona and Bolaina, respectively, to reach enough dry matter (5g) for chemical analysis. In both species, the third and fourth internode from the upper stem were taken as apical samples while the third and fourth internodes from the lower stem as basal samples.

### Wood chemical analysis

The harvested stems were cut into small pieces, dried at 60°C and milled to quantify cellulose, hemicellulose, and lignin content according to Van Soest and Robertson [21]. This method determines neutral detergent fiber (NDF), acid detergent fiber (ADF) and acid detergent lignin (ADL) after digesting samples with chemical reagents. NDF is first obtained after digestion in neutral detergent solution, and it mainly consists of cellulose, hemicellulose and lignin. After digesting NDF with acid detergent, ADF is obtained and contains cellulose and lignin predominantly. To obtain ADL, cellulose was removed from ADF with H_2_SO_4_. Cellulose content is calculated from the difference between ADF and ADL, hemicellulose from the difference between NDF and ADF. Lignin content is expressed as ADL and holocellulose was determined by addition of cellulose with hemicellulose [22]. Analysis of variance (ANOVA) followed by Tukey comparison test (α = 0.05) between the chemical composition of apical and basal stem was performed using R studio.

### FTIR-ATR spectroscopy

Spectra from dried, milled and sieved (60 Mesh) samples of both species were collected. Triplicated FTIR spectra of each sample were acquired by attenuated total reflection (ATR) in Spectrum 100 Perkin Elmer spectrometer equipped with a diamond ATR accessory, a Tantalato de Litio detector and a KBr beam splitter. Thirty-two scans were run per sample in the spectra range between 4000 cm^−1^ and 400 cm^−1^ at 4 cm^−1^ spectral resolution. Before each measurement, background spectra with the same settings were collected.

### Chemometrics

#### Data acquisition and pre-processing

Spectral data were acquired through Spectrum 10 software in SP format. Prior to pre-procesing, spectral data were mean-centered. To remove the effect of interference factors (light scattering, noise, and pathlength variations) from spectral data, the following pre-processing methods (Savitzky-Golay first derivative, Savitzky-Golay second derivative, Smoothing, standard normal variation, multiplicative scatter correction, straight line correction and min-max normalization) were evaluated. Pre-processing was applied in the full spectra region (3700-850 cm^−1^) and the fingerprint region (1800-850 cm^−1^).

#### Partial Least Squares (PLS) Analysis

PLS regression was used to build models for predicting cellulose, hemicellulose, holocellulose and lignin content from FTIR spectra. A cross validation was performed in the Peak spectroscopy software (https://www.essentialftir.com/) to select the best pre-treatment for each compound in the full spectra and the fingerprint region. Once identified the best pre-treatment for each compound, the data were split into calibration set (100 spectra) and validation set (50 spectra). Models were built using calibration set and evaluated using validation set.

#### Determination of number of latent variables

Two methods were performed to determine the number of latent variables: Haaland and Thomas [23] and Gowen et al. [24]. This last method uses the appearance of noise in regression coefficients as a sing of over-fitting. The noise and sharp peaks in regression coefficients were quantified (Jaggedness) and paired to RMSE to estimate the optimal numbers of latent variables [24].

#### Model evaluation

Performance of models were evaluated using root mean squared error of calibration (RMSEC) and predictions (RMSEP), coefficient of determination of calibration 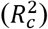 and prediction 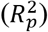, and ratio of performance to deviation (RPD). To assess the predictive ability of models (RPD), Santos et al. [25] and Li et al. [26] formula were used. According to Karlinasari et al [27], models with RPD values higher than 1.5 are satisfactory and useful for initial screenings and preliminary predictions, RPD values between 2.0 and 2.5 have very good prediction and RPD values higher than 3 are efficient prediction. In wood samples an RPD of 1.5 – 2.5 is enough to predict wood chemical composition [28]. Finally, the best model performance was selected based on 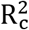 and 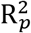 values close to 100, RMSEC and RMSEP values close to cero and a high RPD value [17].

## Results and discussions

### Analysis of chemical composition

The chemical composition of the apical and basal stem for both species (S1 Table) was determined by Van Soest and Robertson [21] method and it is summarized in Table 1. In both species, the percentage of cellulose and holocellulose was significantly higher (p < 0.001) in the basal stem than apical stem. In contrast, a significant decrease in hemicellulose was observed in the basal stem with respect to apical stem in Capirona (p <0.05) and Bolaina (p <0.001). Like carbohydrates, lignin content increased in Capirona basal stem while in Bolaina the lignin content decreased in the apical stem. Therefore, the higher content of carbohydrates and lignin in the base of trees may be related to denser wood as an adaptative response to alleviate any bending stress related to wind and other forces [9,29]. The mechanical benefit of denser wood is the increasing strength in supporting tissues at the base of tree stalk where cellulose provides strength and lignin, compression and shear resistance [30].

**Table 1.** Descriptive statistics of the chemical composition of *C. spruceanum* and *G. crinita*.

| Specie | *Calycophyllum spruceanum* |  |  |  | *Guazuma crinita* |  |  |  |
| --- | --- | --- | --- | --- | --- | --- | --- | --- |
| Compound | % Cel | % Hem | % Hol | % L | % Cel | % Hem | % Hol | % L |
| **Upper steam** |  |  |  |  |  |  |  |  |
| Average | 29.81*** | 10.17* | 39.98*** | 8.14*** | 20.44*** | 19.31*** | 39.75*** | 12.25*** |
| Minimum | 26.24 | 8.08 | 35.84 | 7.05 | 16.48 | 11.56 | 31.54 | 7.82 |
| Maximum | 36.13 | 12.13 | 46.19 | 9.88 | 25.83 | 35.27 | 53.77 | 15.56 |
| SD | 2.47 | 0.92 | 2.47 | 0.64 | 2.03 | 5.92 | 5.48 | 1.89 |
| **Lower steam** |  |  |  |  |  |  |  |  |
| Average | 35.97*** | 9.64* | 45.61*** | 9.65*** | 42.44*** | 14.25*** | 56.69*** | 11.22*** |
| Minimum | 30.91 | 5.49 | 38.99 | 8.14 | 38.83 | 12.42 | 52.04 | 7.26 |
| Maximum | 41.69 | 13.64 | 51.49 | 11.28 | 48.67 | 16.53 | 63.98 | 14.33 |
| SD | 2.94 | 1.39 | 3.16 | 0.67 | 2.04 | 0.86 | 2.37 | 1.35 |

Cel: Cellulose, Hem: Hemicellulose, Hol: Holocellulose and L: lignin
Significance: \* $p < .05$ , \*\* $p < .01$ and \*\*\* $p < .001$ .

### FTIR spectra

In the FTIR spectra of Capirona and Bolaina wood, two regions of analysis were distinguished (3800- 2800 cm-1 and 1800 – 850 cm-1), see Fig. 1. The peak numbers and their assignment to chemical compounds are presented in Table 2. All samples show peaks 1, 2 and 3, which were assigned to O-H stretching, symmetric and asymmetric C-H stretching, respectively (Fig 1 and Table 2) and they are presented in all components of wood [31]. In the fingerprint (1800 – 850 cm-1), there were many peaks assigned to cellulose (peaks 11, 13 and 16), hemicellulose (peak 4), lignin (peaks 6 and 7), holocellulose (peaks 10, 12, 14 and 15) and common peaks to carbohydrates and lignin (peaks 8 and 9).

**Fig 1.**
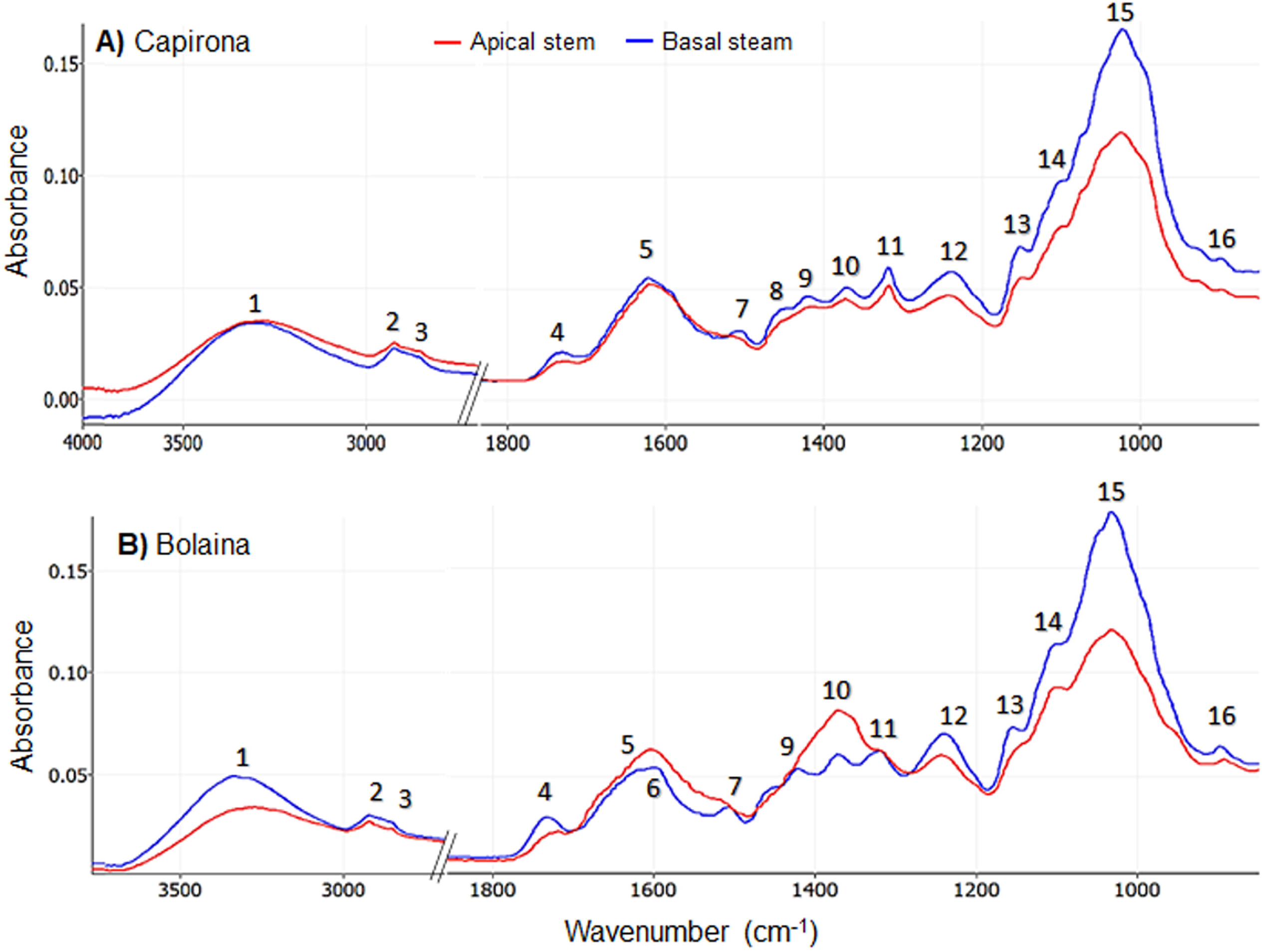
FTIR spectra of juvenile trees. A: Capirona and B: Bolaina.

**Table 2.** Assignment of peaks in FTIR spectra of *C. spruceanum* and *G. crinita*.

| Peak | Boliana |  | Capirona |  | Assignments | References |
| --- | --- | --- | --- | --- | --- | --- |
|  | Upper | Lower | Upper | Lower |  |  |
| 1 | 3266 | 3340 | 3281 | 3315 | Lignin, cellulose and hemicellulose: OH intramolecular and intermolecular stretching. | (32) |
| 2 | 2923 | 2922 | 2923 | 2923 | Lignin, cellulose and hemicellulose: symmetric methyl and methylene stretching. | (32) |
| 3 | 2853 | 2857 | 2854 | 2855 | Lignin and hemicellulose: asymmetric methyl and methylene stretching. | (32) |
| 4 | 1718 | 1733 | 1730 | 1731 | Hemicellulose: C=O stretching in unconjugated ketone, carbonyl, and aliphatic groups. Extractives: C=O stretching in ester carbonyl. | (17,33) |
| 5 | - | 1622 | 1621 | 1623 | Flavones and calcium oxalate: C=O stretching. | (33) |
| 6 | 1603 | 1598 | - | - | Lignin and extractives: aromatic ring vibration. | (17,33) |
| 7 | 1518 | 1508 | 1516 | 1511 | Lignin and extractives: aromatic skeletal vibrations. | (17,33) |
| 8 | - | - | - | 1447 | Cellulose, lignin, and hemicellulose: O-H in-plane bending. | (34) |
| 9 | - | 1422 | 1417 | 1419 | Lignin: C=O stretching and aromatic skeletal vibration. Carbohydrates: CH and OH bending. | (16–18,32) |
| 10 | 1371 | 1372 | 1373 | 1371 | Cellulose and hemicellulose: CH bending. | (32,35) |
| 11 | 1320 | 1319 | 1318 | 1318 | Cellulose: CH<sub>2</sub> wagging. | (32,35) |
| 12 | 1243 | 1240 | 1244 | 1239 | Polysaccharides: C-O stretch and O-H in plane | (36) |
| 13 | - | 1155 | 1150 | 1152 | Cellulose: C–O–C asymmetric stretching | (32) |
| 14 | 1104 | - | - | 1099 | Cellulose and hemicellulose: C–O–C stretching.<br>Lignin: Aromatic C–H in-plane deformation. | (32,35) |
| 15 | 1032 | 1032 | 1024 | 1021 | Holocellulose and lignin: C–O stretching. | (33) |
| 16 | 893 | 897 | 896 | 897 | Cellulose: C1–H deformation | (33) |

Peak 4 corresponds to vibration bonds of C=O in hemicellulose and extractive compounds such as esterified risinic acids, waxes and fats [15,30] and its absorbance was higher in the basal stem than apical stem in both species, which contradicts the chemical analyses (Table 1). These differences could be explained because of the presence of extractive compounds which can be masking hemicellulose vibration [37]. In relation to lignin, peaks 6 and 7 indicate the vibration of the aromatic skeletons [17,33] and there were more intense in the basal stem than apical stem in both species. However, peak 6 was not well defined in Capirona due to the presence of a prominent peak 5, which is attributed to calcium oxalate and flavones [33]. While in Bolaina, peak 5 is poorly defined in apical stem, which could be a signal of a little presence of extractives [34] and it is close to peak 6. Peaks 5, 6 and 7 are in the region from 1750 to 1510 cm^−1^, which was used by Funda et al. [16] to report chemical differences between juvenile and mature wood in Scots pine because of vibrations in lignin and some carbohydrates. Similarly, Ozgenc et al. [38] were able to determine lignin content in alder and cedar based on skeleton of benzenic rings (1508 - 1510 cm^−1^ and 1603 - 1609 cm^−1^). Moreover, peaks 8, 9, 10 and 11 were assigned to stretching and bending bonds of carbohydrates and lignin [9–11,20,22,23]; the absorbance of these peaks in Capirona was higher at basal than apical stem and it was in agreement with chemical analysis (Table 1). For Bolaina, the absorbance of peaks 9, 10 and 11 (Fig. 1B) was higher in the apical section versus basal stem but peak 8 was not distinguished. The spectral region (1510 - 1265 cm^−1^) for peaks 8, 9, 10 and 11 was also used in *Pinus koraiensis* to detect high lignin and carbohydrates concentrations in artificial inclined stem [39]. In the carbohydrate region (1200 to 890 cm^−1^), peaks 12, 13, 14, 15 and 16 (Fig 1A,1B) evidenced a higher concentration of carbohydrates in the basal stem than in the apical stem in both species. Furthermore, within the fingerprint region, Acquah et al. [36] were able to report polysaccharides by the presence of peaks: 897 cm-1, 1030 cm^−1^ (C-H deformation in cellulose and C-O stretch in polysaccharides), 1157 cm^−1^ (C-O-C vibrations) and 1239 cm^−1^ (C-O stretch and O-H stretch in plane in polysaccharides) in forest logging residue. Thus, the FTIR spectra evidenced the compositional difference in different forest samples.

### PLS modeling

PLS regression was applied to develop predictive models that correlate FTIR spectra with chemical wood composition. First, cross validation was performed to select the best pre-treatment for each compound in the full spectra and fingerprint region (S2 Table). The number of latent variables (LV) were determined by Haaland & Thomas [23] method, however higher-order latent variables were observed (S2 Table). Therefore, Gowen et al. [24,40] method was used to prevent overfitting. Then, external validation was performed using the best predictive model obtained by cross validation (Table 3). RMSEC, RMSEP, 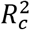, 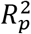 and RPD were used to evaluate the model performance.

**Table 3.** Performance evaluation of PLS models to predict wood chemical composition.

| Species | Component | PP | LV | $R_c^2$ | RMSEC | $R_p^2$ | RMSEP | RPD<sub>a</sub> | RPD<sub>b</sub> |
| --- | --- | --- | --- | --- | --- | --- | --- | --- | --- |
| Capirona | Cellulose | 2<sup>nd</sup>Der | 4 | 85.1 | 1.57 | 74.0 | 2.06 | 1.96 | 1.96 |
|  | Hemicellulose | 1<sup>st</sup>Der | 5 | 77.3 | 0.52 | 61.1 | 0.68 | 1.60 | 1.60 |
|  | Holocellulose | 2<sup>nd</sup>Der | 4 | 83.0 | 1.62 | 72.1 | 2.07 | 1.89 | 1.90 |
|  | Lignin | 2<sup>nd</sup>Der | 4 | 90.2 | 0.31 | 75.4 | 0.48 | 2.02 | 2.06 |
| Bolaina | Cellulose | MM-N | 4 | 97.3 | 1.83 | 97.3 | 1.82 | 6.14 | 6.14 |
|  | Hemicellulose | 1<sup>st</sup>Der | 5 | 85.1 | 1.98 | 81.3 | 2.12 | 2.31 | 2.32 |
|  | Holocellulose | V-N | 6 | 89.3 | 3.09 | 88.8 | 3.16 | 2.99 | 2.99 |
|  | Lignin | 1<sup>st</sup>Der | 6 | 82.6 | 0.72 | 77.7 | 0.81 | 2.12 | 2.12 |

PP: Pre-processing methods: (1<sup>st</sup> Der: First derivative, 2<sup>nd</sup> Der: Second derivative. MM-N: Min - Max normalization and V-N: Vector normalization).
LV: Latent variables according to Gowen et al. [21].
<sup>a</sup> RPD according to Santos et al. [25].
<sup>b</sup> RPD according to Li et al. [26].

The lignin models of Capirona and Bolaina (Table 3) were highly accurate because of low RMSEP values (0.48 and 0.81, respectively), with a good prediction power (RPD > 2.0) and a good data fitting (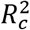 of 90.2% and 82.6%). Our results are in agreement with Acquah et al. [36], Zhou et al. [17], Zhou et al. [37], who reported values of R^2^ ≈ 90 – 80 % and RMSEP ≈ 0.47 - 0.80. Despite these authors reported models with lower RPD (1.4 and 1.72) values than our models’ (see table 3), the author models were considered to have a good predictive power.

In Capirona, the predictive power of cellulose (Fig. 2A), hemicellulose (Fig. 2B) and holocellulose (Fig. 2C) models were satisfactory to quantify these compounds. The RPD values of these three compounds were within 1.5 to 2.0 range (Table 3), they are suitable for initial screenings and preliminary predictions [26,27]. However, these models would be proper for estimating wood properties according to Hein et al. [28]. Moreover, the 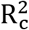 values for cellulose and holocellulose content were 85.1% and 83.0 %with a RMSEP of 2.06 and 2.07, respectively. Meanwhile, the 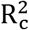 value for hemicellulose was 77.3% with a RMSEP of 0.68. A good hemicellulose predictive model with similar values to our model was reported by Funda et al. [18] in juvenile wood of Scots pine. Furthermore, our cellulose model is consistent with Zhou et al. [17], who reported R^2^ and RPD values of 85.62% and 1.72 in hardwoods so this model was strong enough for screening and cellulose characterization. Finally, our holocellulose model was better than the one reported by Acquah et al. [36] in forest logging residue. On the other hand, Funda et al. [18], Zhou et al. [17] and Acquah et al. [36] reported more efficient lignin models based on a high RPD and a low RMSEP than carbohydrate models. We found this trend in Capirona models too. A possible explanation to this trend may be the presence of unique chemical structures in lignin unlike carbohydrates [36].

**Fig 2.**
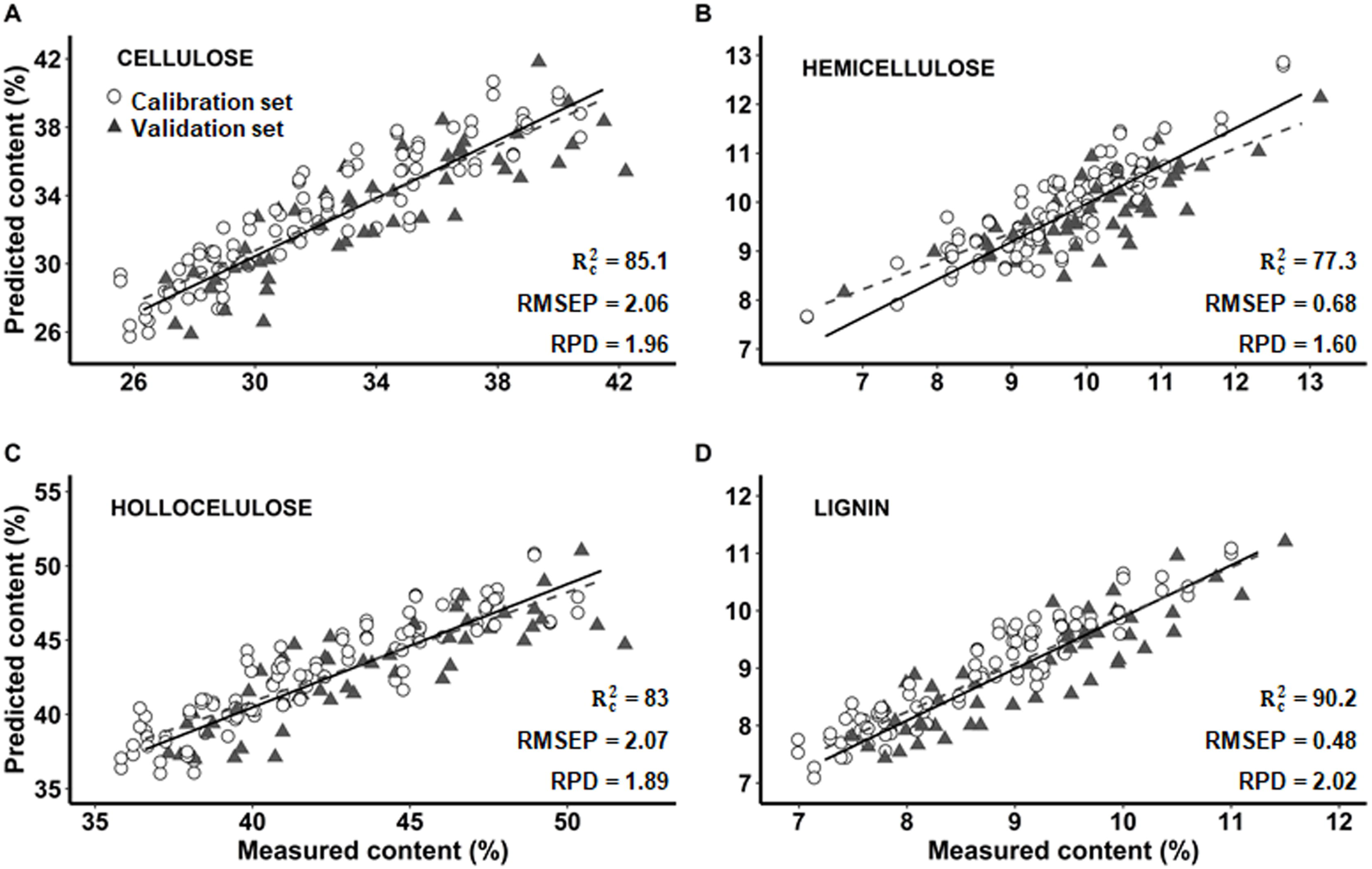
Prediction of cellulose, hemicellulose, hollocelulose and lignin content by multivariate models with FT-IR (PLS) in Capirona (*C. spruceanum*). A: Cellulose, B: Hemicellulose, C: Holocellulose and D: Lignin. 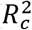: coefficient of determination of calibration. RMSEP: root mean squared error of predictions. RPD: ratio of performance to deviation.

In Bolaina, the RMSEP values of cellulose, hemicellulose and holocellulose (1.82, 2.12 and 3.16, respectively) were higher compared to other authors. For instance, Funda et al. [18], who reported RMSEP values from 0.69 to 0.71 in cellulose and 0.71 in hemicellulose, while Zhou et al. [17] found RMSEP values from 0.8 to 1.9 in cellulose and hemicellulose. Since R^2^ value depends on the distribution of the data and not only on the model performance (41), it is possible to use it as a secondary measure of model performance [18]. Thus, the performance models should achieve RMSEC and RMSEP values close to zero, a high RPD with 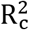 and 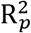 values close to 100. In table 3, models showed a high predictive power (RPD > 2) with RMSEP slightly high, so R2 was considered to evaluated model performance. Thus hemicellulose (Fig 3B) and holocellulose (Fig 3C) models had a high data correlation (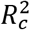 of 85.1 and 89.3, respectively) and RPD > 2.0, so they were considered as very good prediction models [26,27]. Cellulose model (Fig 3A) showed exceptionally high values of 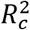=97.3 and RPD = 6.14, so it was classify as an excellent predictive model [26,27]. In contrast, the cellulose, hemicellulose and holocellulose models reported by Acquah et al. [36] in forest logging residue and Zhou et al. [17] in hardwoods were considered satisfactory.

**Fig 3.**
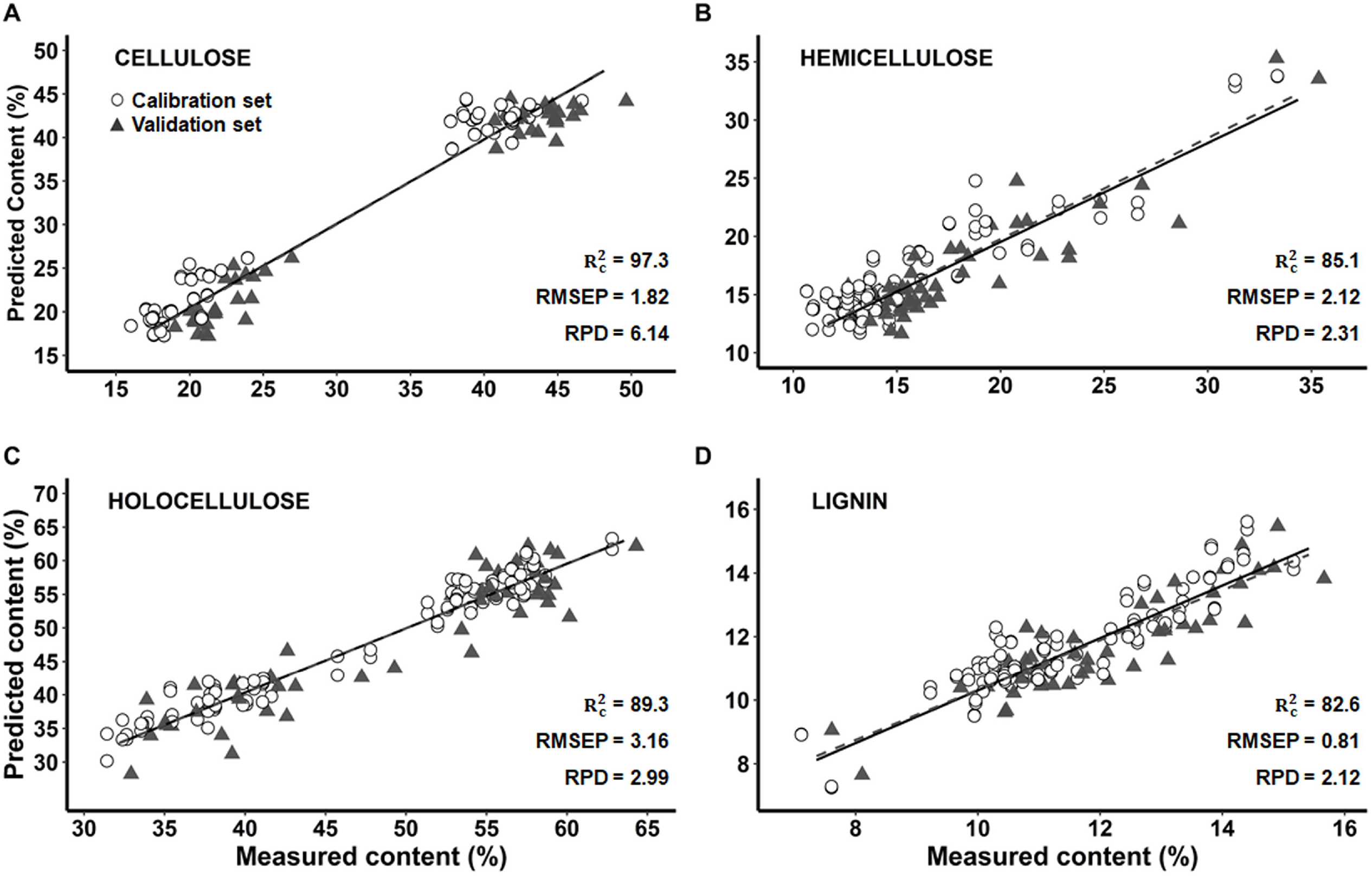
Prediction of cellulose, hemicellulose, hollocelulose and lignin content by multivariate models with FT-IR (PLS) in Bolaina (*G. crinita*). A: Cellulose, B: Hemicellulose, C: Holocellulose and D: Lignin. 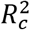: coefficient of determination of calibration. RMSEP: root mean squared error of predictions. RPD: ratio of performance to deviation.

On the other hand, in Bolaina lignin model (Fig. 3D) had a higher accuracy (RMSEP = 0.81) than carbohydrates models (RMSEP from 1.82 to 3.16), which agrees with *Pinus sylvestris* [18], forest logging residue [36] and hardwoods [17] models. However, the lignin model presented lower predictive power (RPD = 2.12) than the carbohydrates models (RPD from 2.32 to 6.14). In NIR, Karlinasari et al. [27] and Jiang et al. [22] reported this trend in *Acacia mangium* and *Pinus spp*, respectively.

For Capirona and Bolaina, the accuracy of the models (RMSEP) was higher in lignin than the carbohydrate models (Table 2). According to Hein et al. [25], an RPD > 1.5 allows to quantify the chemical composition of the wood. However, the RPD scale established by Karlinasari et al. [27] allows a better classification of the models. Finally, the values of R^2^ and RMSE were close between the set of our calibration and validation models, therefore models have good predictive capacity according to Li et al. [26].

## Conclusions

A rapid method for wood chemical determination in an integrated production system based on Capirona (international market) and Bolaina (national marker) is required for genetic improvement and hence impulse a forest-based bioeconomy in Amazon. This study confirmed significant differences in cellulose, hemicellulose, holocellulose and lignin content between apical and basal stem in Capirona and Bolaina. The variability found was used to build PLS models to estimate cellulose, hemicellulose, holocellulose and lignin content in juvenile wood from FTIR-ATR spectra. Thus, lignin models of both species achieved high accuracy (low RMSEP) and were considered as very good prediction models (RPD > 2). In Bolaina, models of hemicellulose and holocellulose showed a very good prediction power (RPD > 2.0), meanwhile cellulose model was excellent (RPD > 3.0). In Capirona, cellulose, hemicellulose and holocellulose models attained sufficient predictive power (RPD > 1.5) to estimate wood properties. Thus FTIR-ATR spectroscopy combined with PLS models offers a technologic tool applied to early tree selection based on chemical wood composition.

## Acknowledgment

The authors would also like to thank Vladimir Camel for his critical reading.

## Supporting Information

**S1 Table.** Wood chemical composition

**S2 Table.** Leave-one-out cross validation method and number of latent variables determination

